# Some theoretical aspects of reprogramming the standard genetic code

**DOI:** 10.1101/2020.09.12.294553

**Authors:** Kuba Nowak, Paweł Błażej, Małgorzata Wnetrzak, Dorota Mackiewicz, Paweł Mackiewicz

## Abstract

Reprogramming of the standard genetic code in order to include non-canonical amino acids (ncAAs) opens a new perspective in medicine, industry and biotechnology. There are several methods of engineering the code, which allow us for storing new genetic information in DNA sequences and transmitting it into the protein world. Here, we investigate the problem of optimal genetic code extension from theoretical perspective. We assume that the new coding system should encode both canonical and new ncAAs using 64 classical codons. What is more, the extended genetic code should be robust to point nucleotide mutation and minimize the possibility of reversion from new to old information. In order to do so, we follow graph theory to study the properties of optimal codon sets, which can encode 20 canonical amino acids and stop coding signal. Finally, we describe the set of vacant codons that could be assigned to new amino acids. Moreover, we discuss the optimal number of the newly incorporated ncAAs and also the optimal size of codon blocks that are assigned to ncAAs.

## 2 Introduction

The standard genetic code (SGC) is a set of rules according to which 64 codons are assigned to 20 canonical amino acids and stop coding signal. Thanks to that, genetic information can be stored in DNA and transmitted into the protein world. It is clear that the SGC is redundant because there are 18 amino acids encoded by more than one codon, i.e. 2, 3, 4 and 6 codon blocks or boxes. Therefore, it seems reasonable to reduce this redundancy in order to extended the genetic code. Thanks to that, we could use the spare codons for introducing new genetic information into the canonical coding system. The inclusion of non-canonical amino acids (ncAAs) in the code can allow us for production of new artificial proteins with novel functionality. This approach is very promising for synthetic biology and can find many applications in medicine, industry and biotechnology.

There are several approaches to the problem of the SGC extension [Chin, 2014]. The first one is stop-codon suppression [Noren et al., 1989, Chin, 2017, Italia et al., 2017, Young and Schultz, 2018]. In this method, stop translation codons, for example *U AG*, are used to encode new ncAAs. This technique needs a modified aminoacyl-tRNA synthetase that charges a tRNA molecule with the ncAA. However, this approach has several drawbacks. For example, we can extend the SGC by only up two new amino acids, because one of the three stop codons must be left to act as a termination signal of translation [Ozer et al., 2017]. What is more, the newly added ncAAs could compete with translation release factors, which may have an impact on the quality of the protein synthesis. The second method is related to programmed frameshift suppression. In this approach, four-base codons (quadruplets) are used to incorporate new ncAAs [Hohsaka et al., 1996, Anderson et al., 2004, Neumann et al., 2010]. Generally, these codons are composed of a rarely used classical codons with an additional base. These structures are decoded by a modified tRNAs containing four-base anticodons. It should be noted that the competition between tRNAs reading classical codons and respective quadruplet codons can decrease the efficiency of the whole procedure. The third method postulates the extension of the standard genetic code by using selected synonymous codons and depletion of their corresponding tRNAs, which are pre-charged with ncAAs [Iwane et al., 2016]. This method enables significant increasing the number of ncAAs by using of multiple codon boxes. However, the using of the current synonymous codons can disturb the translation and protein folding because of change in codon usage and speed of protein synthesis [Plotkin and Kudla, 2011]. Other approach is based on using one pair of unnatural nucleotides [Ishikawa et al., 2000,Ohtsuki et al., 2001, Yang et al., 2007,Kimoto et al., 2009, Malyshev et al., 2009, Dien et al., 2018,Hamashima et al., 2018]. Thereby it is possible to generate up to 152 new codons to which ncAAs can be assigned. Thanks to that, new genetic information does not interfere with the natural system because it does not involve the canonical codons.

A theoretical approach in the extension of the SGC has been recently proposed by [Błażej et al., 2020]. The authors analysed how to extend the SGC up to 216 codons generated by six-letter nucleotide alphabet, including besides four canonical bases also one pair of new bases. The model of the code assumes the gradual addition of the codons to minimize the consequences of point mutations. In this paper, we would like to investigate other theoretical aspects of the standard genetic code extension using 64 canonical codons. Following the redundancy of the SGC, we focused on finding the rules of the SGC expanding via optimal partition of codon boxes. In our first step, basing on the methodology incorporated from graph theory, we found the minimal set of codons that encode the whole canonical information that is at the same time the most robust to lose this information via point mutations. Therefore, we could divide the set of all 64 codons into two subsets: the first one encoding classical repertoire of amino acids and the second one that is composed of potentially vacant codons. Moreover, we studied some scenarios of optimal codon reassignments that cause creating a new extended coding system.

## 3 Preliminaries

We would like to discuss some properties of the SGC using the methodology of graph theory. This approach was successfully used in many problems related to the standard genetic code optimality, in which the SGC is represented as a partition of selected graph [Błażej et al., 2018a, Błażej et al., 2019b]. Some rules of the optimal genetic code enlargement using the extended nucleotide alphabet were presented [Błażej et al., 2020].

We started our considerations with describing the graph that represents relationships between all possible 64 codons in terms of point mutations. Similarly to [Błażej et al., 2018a, Błażej et al., 2019b, Błażej et al., 2020], let *G*(*V, E*) be a graph, in which *V* is the set of vertices representing all possible 64 codons, whereas *E* is the set of edges between these vertices. We say that two codons *u, v* ∈ *V* are connected by the edge *e*(*u, v*) ∈ *E* if and only if the codon *u* differs from the codon *v* in exactly one position. The graphical representation of *G* is given in the Figure 1.

**Figure 1.**
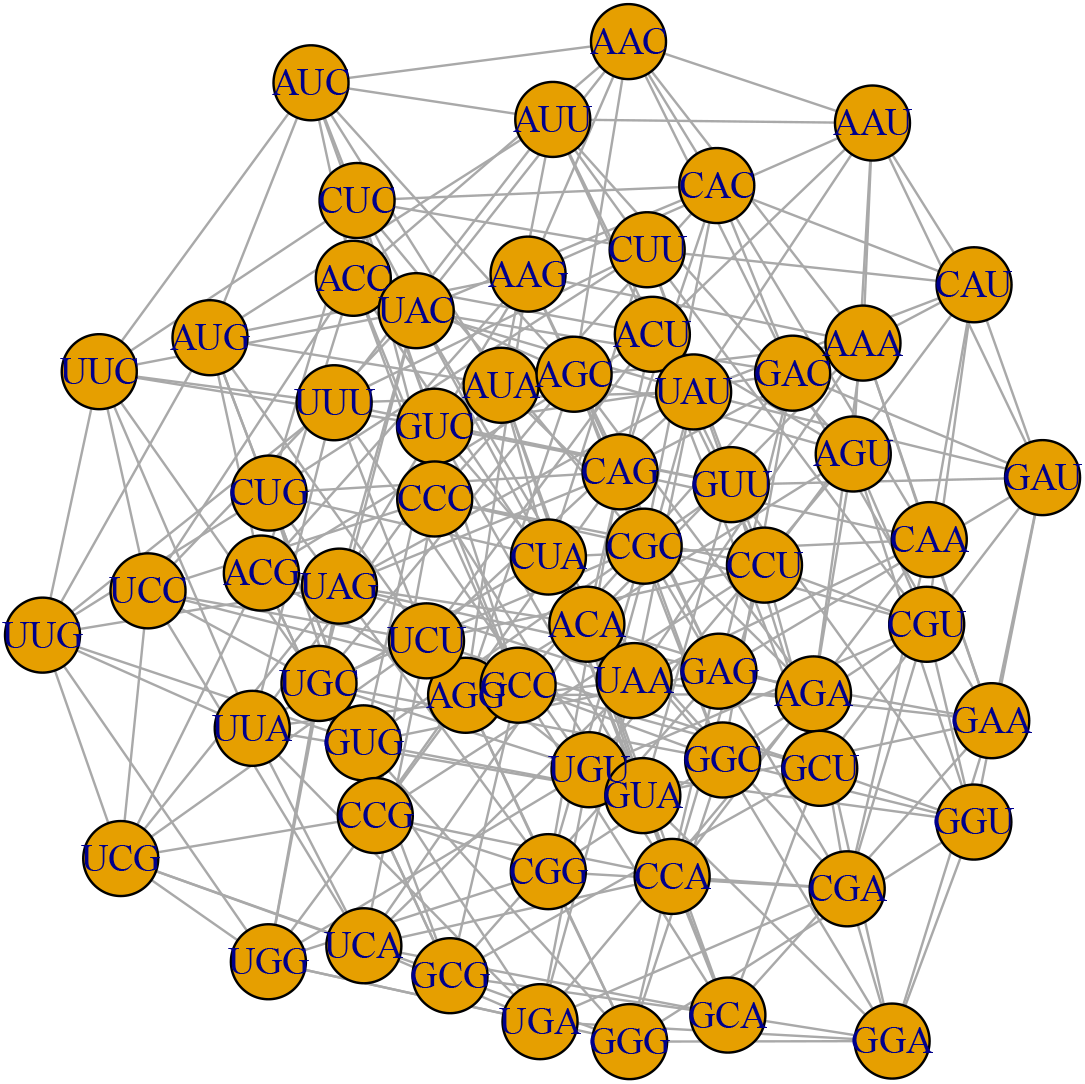
The representation of the graph *G*(*V, E*), in which the set of all vertices *V* is composed of 64 codons, whereas the set of edges *E* is induced by all possible point mutations that might occur in codons. Clearly, *G* is regular and the degree of each vertex is equal to nine. We do not take into account any additional properties of the mutational process, hereby *G* is also unweighted and undirected.

Clearly, *G* is an unweighted and regular graph. Interestingly, *E* represents all possible single point mutations, which may occur between codons in protein coding sequences. It should be noted that here we do not take into account any additional properties of the mutational process, e.g. different probability (weight) of individual nucleotide substitutions.

Following graph theory, every possible genetic code, including its extension, induces a partition 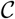 of the set *V* into *l* ≥ 21 disjoint non-empty subsets *S*. Therefore, we can introduce the following, unique representation:

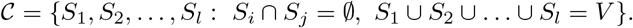

In the papers [Błażej et al., 2018a, Błażej et al., 2019b, Błażej et al., 2020], the authors investigated the properties of the optimal partition of the graph *G* in terms of the set conductance. Here, we apply a similar characteristics, i.e. the *k*-size conductance and the average conductance of set collection, which were defined below.

### Definition 1.

*For a given graph G, let S be a subset of V. The conductance of S is defined as:*

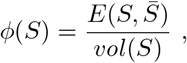

*where* 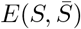 *is the number of edges of G crossing from S to its complement* 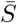 *and vol(S) is the sum of all degrees of the vertices belonging to S*.

The set conductance *ϕ*(*S*) has an interesting interpretation. Let us assume that *S* is a codon block encoding a selected amino acid. Then, *ϕ*(*S*) give us an information about the robustness level of this amino acid to point mutations that change it other amino acid. In other words, it is a measure of losing information in the system. This observation rises immediately the question about the minimum set conductance for sets with a given size *k*. It is particularly interesting in the context of the optimal encoding of genetic information by a block of codons. In order to study this property, we used the following definition.

### Definition 2.

*The k-size-conductance of the graph G, for k* ≥ 1, *is defined as:*

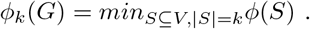

The last characteristic, called the average conductance of set collection, allow us to evaluate the general quality of genetic code under study.

Let us define the average conductance.

### Definition 3.

*Let* 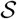 *be a set collection that fulfils the following property:*

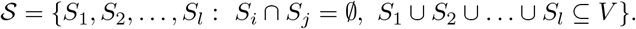

*The average conductance of* 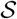 *is defined as:*

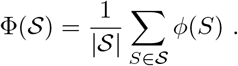

It is easy to notice that 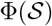 is a generalisation of the average code conductance presented in [Błażej et al., 2018a]. In fact, if 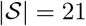 and *S*_1_ ∪ *S*_2_ ∪ … ∪ *S_l_* = *V* are codon blocks created according to the SGC rules, then 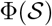 is the average code conductance for the standard genetic code. Clearly, the average conductance of set collection gives us a general information about the properties of all codon blocks *S_i_, i* = 1, 2,…, *l* which constitute 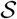.

These characteristics appeared to be very useful in studying the structural properties of genetic codes. However, they require a fast and effective method for determining the optimal *ϕ_k_*(*G*) for *k* ≥ 1. Fortunately, the graph *G* possesses many interesting properties, which were discussed in [Błażej et al., 2018a, Błażej et al., 2020]. First of all, *G* can be represented as a Cartesian graph product, i.e.

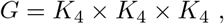

where *K*_4_ is a 4-clique with the set of vertices corresponding to nucleotides {*A, U, G, C*}. This property allows us to characterise the set of vertices reaching the minimal set conductance from all possible subsets with a given size *k*. The following proposition presented in [Błażej et al., 2018a] is a natural consequence of the Theorem 1 given by [Bezrukov, 1999].

### Proposition 1.

*Let us consider a linear order of the set of vertices of* 4-*clique K*_4_, *for example A > C > G > U, and let O_k_ be a collection of the first k vertices of a graph K*_4_ × *K*_4_ × *K*_4_ = *G in the lexicographic order. Then we get:*

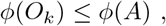

where *A* ⊆ *K*_4_ × *K*_4_ × *K*_4_, |*A*| = *k, for any k* ≥ 1. *Therefore, the following equations hold for any k* ≥ 1:

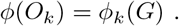

As a result, each sequence of *k*, 1 < *k* < 64 vertices of *G* sorted according to a given lexicographic order can reach the minimum of the set conductance over all possible set of vertices with the size *k*. We used the notation *O_k_* in the whole paper to denote the general set of codons in the lexicographic order. We described the order of codons when it was necessary (Tab. 1). Thanks to that, we avoided a complicated and redundant notation.

**Table 1.** The example of the set *O_k_* for *k* = 8, which is a sequence of the first eight codons taken in a selected lexicographic order. According to Proposition 1 this set is characteriZed by the minimal set conductance over all sets with the size of *k* = 8. The codons have assigned encoded amino acids as in the standard genetic code.

| codon | amino acid |
| --- | --- |
| AAA | Lys |
| AAC | Asn |
| AAG | Lys |
| AAU | Asn |
| ACA | Thr |
| ACC | Thr |
| ACG | Thr |
| ACU | Thr |

It should be noted that including all possible linear orders of the set {*A, U, G, C*} and also all possible orders of codon positions {1, 2, 3}, there are exactly 144 different lexicographic orders which can be introduced to *G*.

## 4 Results and discussion

We began our investigation with describing the smallest set of codons encoding all 20 amino acids and stop coding signal, which still preserves the canonical codon assignments and is simultaneously optimal in terms of the set conductance *ϕ*. Then, using the complement set of vacant codons, we discussed some scenarios of reprogramming the standard genetic code. Particularly, we would like to investigate properties of some extensions of the SGC under the less restrictive assumptions. In this case, we used the average conductance of the set collection Φ as a measure of the quality of a given genetic structures, i.e. codon blocks. Finally, we studied the specific way of code reprogramming that includes the general structure of the standard genetic code.

The requirements imposed on the genetic code in this approach assume its robustness to changes causing the loss of genetic information. This assumption follows the adaptation hypothesis, which claims that the SGC evolved to minimize harmful consequences of mutations or mistranslations of coded proteins [Woese, 1965, Sonneborn, 1965, Epstein, 1966, Goldberg and Wittes, 1966, Haig and Hurst, 1991, Freeland and Hurst, 1998, Freeland et al., 2000,Gilis et al., 2001]. Although this code did not turn out perfectly optimized in this respect [Błażej et al., 2018a, Błażej et al., 2016, Massey, 2008, Novozhilov et al., 2007, Santos et al., 2011, Santos and Monteagudo, 2017, Wnetrzak et al., 2018, Błażej et al., 2018b, Błażej et al., 2019b, Wnetrzak et al., 2019], it shows a general tendency to error minimization in the global scale. This property is better exhibited by its alternative versions [Błażej et al., 2018c, Błażej et al., 2019a], which occurred later in the evolution. Therefore, the analysis of the genetic code extension in this context seems to be a natural consequence of its evolution.

### 4.1 The smallest set of codons encoding canonical information

It is well known that the standard genetic code is redundant, which means that a smaller number of codons is enough to encode all 20 canonical amino acids and one stop translation signal. Theoretically, we can randomly select more than 1.51 * 10^80^ genetic codes but many of them will not effectively function in biological systems, which can require all 20 canonical amino acids and one stop translation signal as well as some specific relationships between codons. In the selection, we assume that the chosen codons have amino acid assignments as in the SGC. Therefore, it seems reasonable to postulate some conditions that must be met by these minimalistic genetic codes. In this work, we assume that the codon sets must fulfill some properties in terms of the set conductance φ. This assumption has a sensible biological meaning because this measure represents the ratio of non-synonymous mutations, which change a given amino acid or stop codon to other, to all possible point mutations. The small value of *ϕ* indicates that the code is resistant to change in the coded information.

Following Proposition 1, we get that the first *k*-codons ordered in lexicographic order *O_k_* constitute the set with the minimum set conductance φ over all possible sets of codons with the size *k* (Tab. 1). In consequence, it is the most resistant structure against loosing information stored in this set for 1 < *k* < 64. This property poses a question about the minimum number of codons *k* such that there exists a set *O_k_* composed of codons that encode 20 amino acids and stop translation signal. In order to deal with this problem, we denote *C_k_* as the set of *k* lexicographically ordered codons that encode 21 canonical items. Moreover, for the selected set *C_k_*, we define a collection of sets:

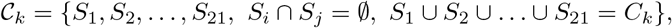

where *S_i_, l* = 1, 2,…, 21 is a non-empty set of codons encoding genetic information according to the standard genetic code rules. Here, we tested all possible *C_k_* induced by all 144 orders. As a result, we obtained that *k* = 28 is the minimal siZe of codons set for which there exists a well defined *C_k_*, i.e. codons belonging to *C*_28_ encode 20 amino acids and stop coding signal. In fact there are two lexicographic orders, the first is induced by a linear order between nucleotides *U* < *G* < *A* < *C* and an order between codon positions 1 < 2 < 3. The second is generated by a linear order *G* < *U* < *A* < *C* between nucleotides and an order between codon positions 1 < 2 < 3. We found that the first 28 ordered codons using these two rules encode all 21 items. Table 2 and Table 3 include representations of 64 codons in the classical standard genetic code table showing the structure of the optimal *C*_28_ codon set. All codons belonging to *C_k_* are marked in red. We also presented the names of encoded items, which induces at the same time the set collection 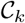.

**Table 2.** The set *C*_28_ containing exactly 28 codons (red). These codons were chosen according to a lexicographic order induced by the linear order of nucleotides *U* < *C* < *A* < *G* and the order of codon positions 1 < 2 < 3. The encoded 20 canonical amino acids and the stop translation signal by the collection of codon sets 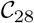 is also shown.

|  |  |  |  |
| --- | --- | --- | --- |
| UUU Phe | UCU | UAU Tyr | UGU Cys |
| UUC | UCC | UAC | UGC |
| UUA | UCA | UAA | UGA |
| UUG Leu | UCG Ser | UAG Stop | UGG Trp |
| CUU Leu | CCU | CAU His | CGU Arg |
| CUC | CCC | CAC | CGC |
| CUA | CCA | CAA | CGA |
| CUG Leu | CCG Pro | CAG Gln | CGG Arg |
| AUU Ile | ACU | AAU Asn | AGU Ser |
| AUC | ACC | AAC | AGC |
| AUA | ACA | AAA | AGA |
| AUG Met | ACG Thr | AAG Lys | AGG Arg |
| GUU Val | GCU | GAU Asp | GGU Gly |
| GUC | GCC | GAC | GGC |
| GUA | GCA | GAA | GGA |
| GUG Val | GCG Ala | GAG Glu | GGG Gly |

**Table 3.** The set *C*_28_ containing exactly 28 codons (red). These codons were chosen according to a lexicographic order induced by the linear order of nucleotides *G* < *U* < *A* < *C* and the order of codon positions 1 < 2 < 3. The encoded 20 canonical amino acids and the stop translation signal by the collection of codon sets 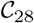 is also shown.

|  |  |  |  |
| --- | --- | --- | --- |
| UUU Phe | UCU Ser | UAU Tyr | UGU Cys |
| UUC | UCC | UAC | UGC |
| UUA | UCA | UAA | UGA |
| UUG Leu | UCG | UAG Stop | UGG Trp |
| CUU Leu | CCU Pro | CAU His | CGU Arg |
| CUC | CCC | CAC | CGC |
| CUA | CCA | CAA | CGA |
| CUG Leu | CCG | CAG Gln | CGG Arg |
| AUU Ile | ACU Thr | AAU Asn | AGU Ser |
| AUC | ACC | AAC | AGC |
| AUA | ACA | AAA | AGA |
| AUG Met | ACG | AAG Lys | AGG Arg |
| GUU Val | GCU Ala | GAU Asp | GGU Gly |
| GUC | GCC | GAC | GGC |
| GUA | GCA | GAA | GGA |
| GUG Val | GCG | GAG Glu | GGG Gly |

In the next step, we test the quality of 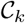, *k* = 28,…, 63 structure in terms of the average conductance of set collection Φ. It is clear that 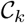 is generated by the lexicographically ordered set of codons and the canonical codon assignments. Using Φ, we can compare different genetic code structures generated by different lexicographic orders for the same number of codons *k*. Figure 2 presents a relationship between 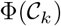 and *k* = 28, 29,…, 64 calculated for two lexicographic orders, for which we found the smallest coding set *C*_28_ (blue and orange lines). The lower bound calculated over 144 orders is shown for comparison (green line). As we can see, in all considered cases Φ decreases with the number of codons involved in the set. They all reach the maximum at *k* = 28, which is equal 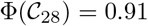, and the minimum for all set collections at *k* = 64, equal to 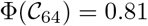. It should be noted that 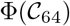 is equal to the average code code conductance calculated for the standard genetic code and discussed in [Błażej et al., 2018a]. What is more, two lexicographic orders that generate the respective smallest codon sets *C*_28_, generally do not induce the optimal collections of sets 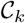, *k* > 28 in terms of Φ. In other words, it is not possible to generate a set collection 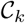 for each *k* = 29,… 64 using lexicographic orders shown in Table 2 and Table 3 that would be minimal in terms of Φ.

**Figure 2.**
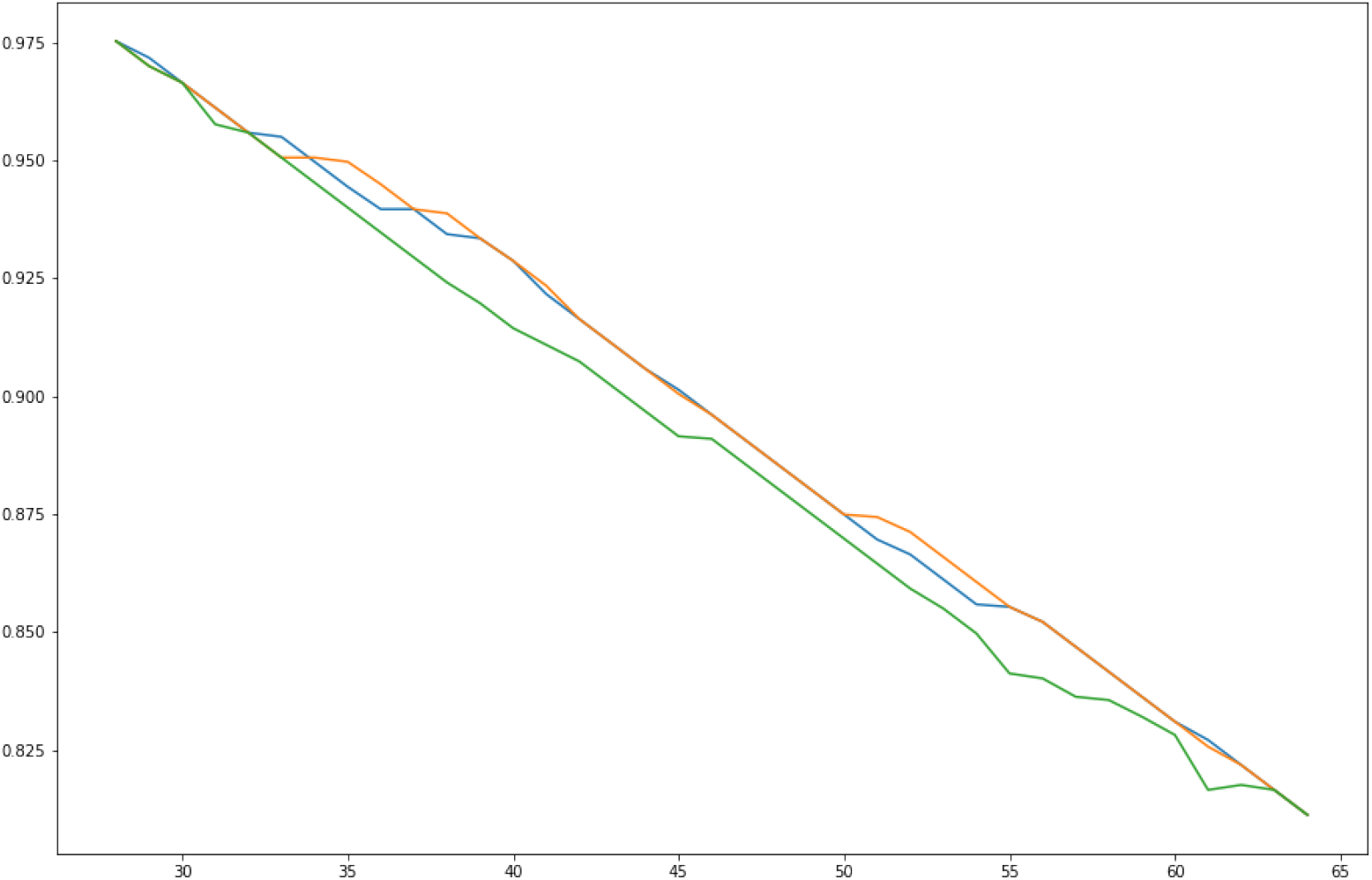
The relationship between 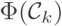 and *k* = 28, 29,…, 64 calculated for two lexicographic orders, for which we found the smallest coding set *C*_28_ (blue and orange lines). The lower bound calculated over 144 orders is shown for comparison (green line).

### 4.2 Reprogramming of the standard genetic code

Using the results presented in section 4.1, we get that for every *k* ≥ 28, the set *C_k_* induces its own complement, namely the set 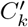 of vacant codons. They can be reprogrammed to encode new non-canonical amino acids. From mathematical perspective, new genetic information would be encoded by 1 ≤ *n* ≤ 64 – *k* codon blocks which constitute a partition of the set 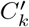. In consequence, we introduced a set collection of *n* codon blocks:

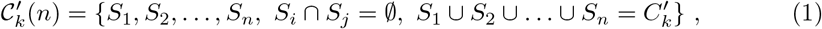

where each *S_i_, i* = 1,… *n* is a non-empty set of codons that encodes the same genetic information. Moreover, let us denote by 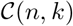 an extended genetic code that encodes exactly *n* new ncAAs and uses a set *C_k_, k* ≥ 28 to maintain canonical genetic information. It is easy to see that every 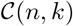 is a partition of the graph *G*. What is more, 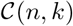 has a unique representation:

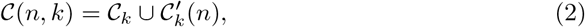

for any *k* ≥ 28. This observation appeared to be very useful in a further testing of the properties of the extended genetic code. First of all, it is possible to calculate 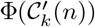, i.e. the average conductance of the set collection encoding ncAAs.

Additionally, 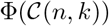 gives us a general overview on the properties of the extended genetic code. Furthermore, using equation 2 we can measure the structural differences between the canonical 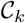 and the extended 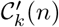 set collection. In order to do so, we introduced a balance measure *B* defined in the following way:

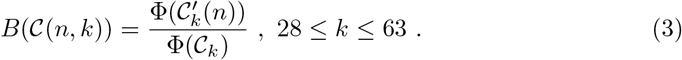

The balance function *B* has a natural interpretation and takes positive values. *B* < 1 indicates that 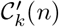 possesses better structural properties in terms of the average conductance than 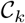, whereas *B* > 1 means that the set collection 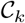, i.e. canonical genetic information has better properties in terms of Φ. From our point of view, the value of B around one is the most interesting because it suggests a similar robustness to point mutations of codon blocks for both set collections.

#### 4.2.1 The optimal codon blocks structures encoding ncAAs

Here, we discuss several features of set 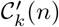, *k* ≥ 28, *n* = 1,…, 64 – *k*, which is a collection of *n* codon blocks composed of vacant codons 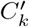. The problem which arose immediately in this study was related to the potential optimality of 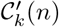. As was mentioned in the previous section, we can describe the quality of each 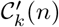 using the average conductance of the set collection Φ. However, this measure itself does not give us any information about that the real optimality of a studied code because there are, in general, many possible set collections for the fixed *k* and *n* that differ in Φ values. Therefore, we decided to find a lower bound on the values of 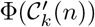 for the fixed *k* ≥ 28 and *n* = 1,… 64 – *k*. It could be done using the representation 1 of 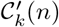 as well as the definition 2 of the *k*-size conductance and a simple observation that for every 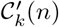, we have:

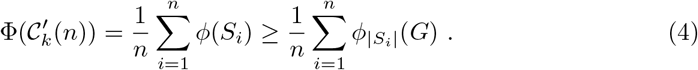

Therefore, for every 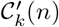, there exists a lower bound on the average conductance of the set collection imposed on 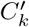. What is more, these optimal collections are composed of the best codon blocks in terms of the k-size conductance. This feature give us a general overview on the optimal structures of the standard genetic code extensions including the selected number *n* of ncAAs.

Following the property 4, we found all possible lower bounds for every *k* ≥ 28 and *n* = 1, 2,… 64 – *k*. Figure 3 presents their graphical representations. As we can see, the lower bound on 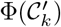 increases with the partition size n for all considered *k* ≥ 28.

**Figure 3.**
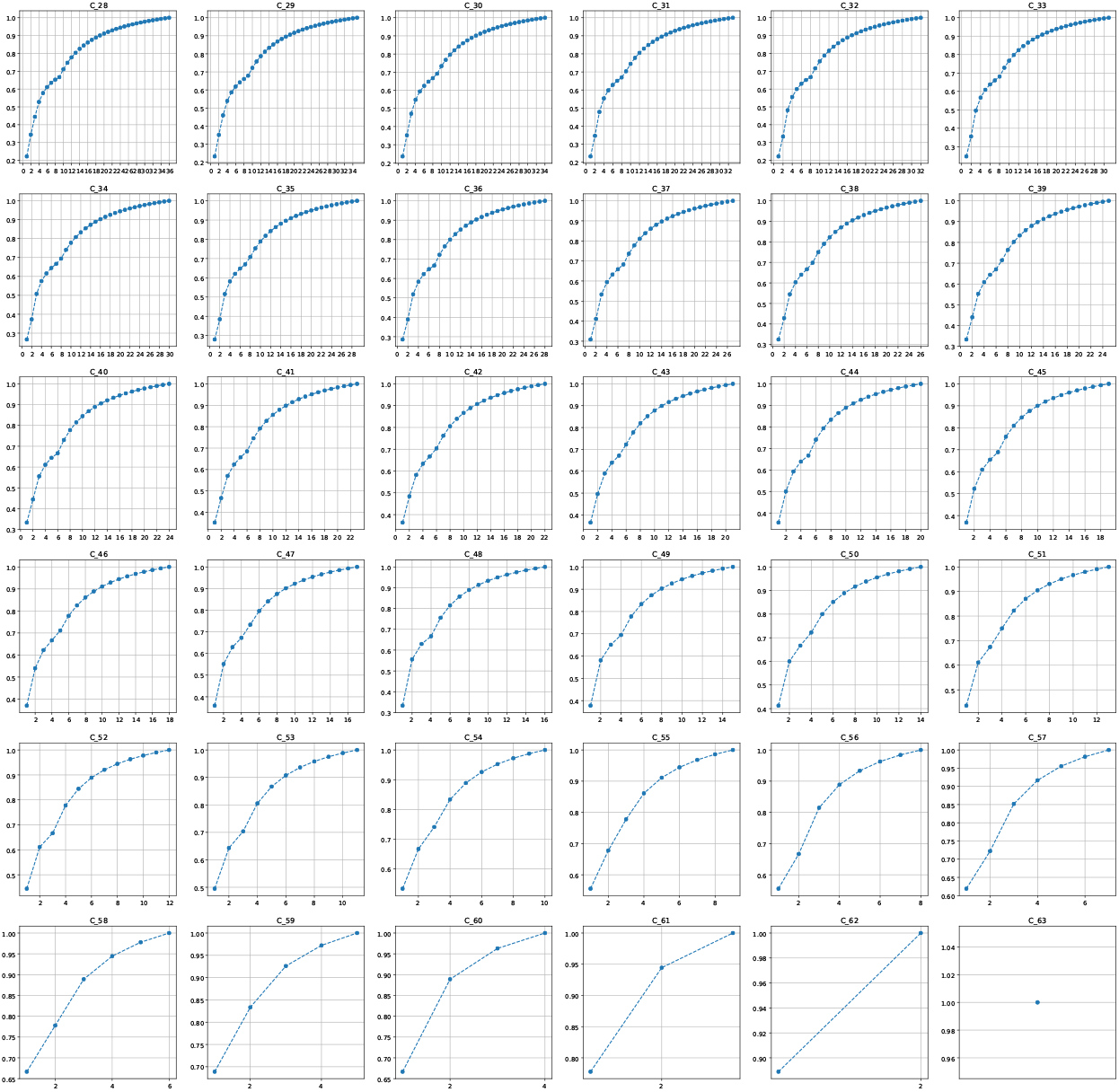
The lower bound of the average conductance 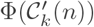 calculated for the set collection 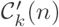 in relation to the number *n* of potential codon blocks which would encode new genetic information. The minimum of 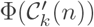 was found over all possible partitions of the set 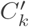 containing *k* ≥ 28 vacant codons in exactly *n* = 1,…, 64 – *k* disjoint codon blocks.

This relationship shows an interesting course in some cases, e.g. for *k* = 28 (Fig. 4), the curve of the lower bound increases with *n* but slows down substantially for *n* close to *n′* = 9 and then blow up again for *n* > *n′*. We can explain this fact by comparing the properties of the optimal 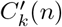 partition for *n* = 1, 2,…, 36. We realized that for *n* ≤ *n′*, there is no set of the size lower than four, whereas these sets appear for *n* > *n′*. It should be noted that the *k*-size conductance is decreasing with *k*, especially for *k* = 1, 2, 3. Therefore, if a set collection contains a group of size that is lower than four then the average conductance of the set collection calculated for this partition is generally higher in comparison to the collections that are composed of codon blocks with the size greater or equal than four. This fact could explain the presence of the Φ minimum for *n* > *n′*. This phenomenon is also observed for 28 ≤ *k* ≤ 52 for respective changing point *n′*.

**Figure 4.**
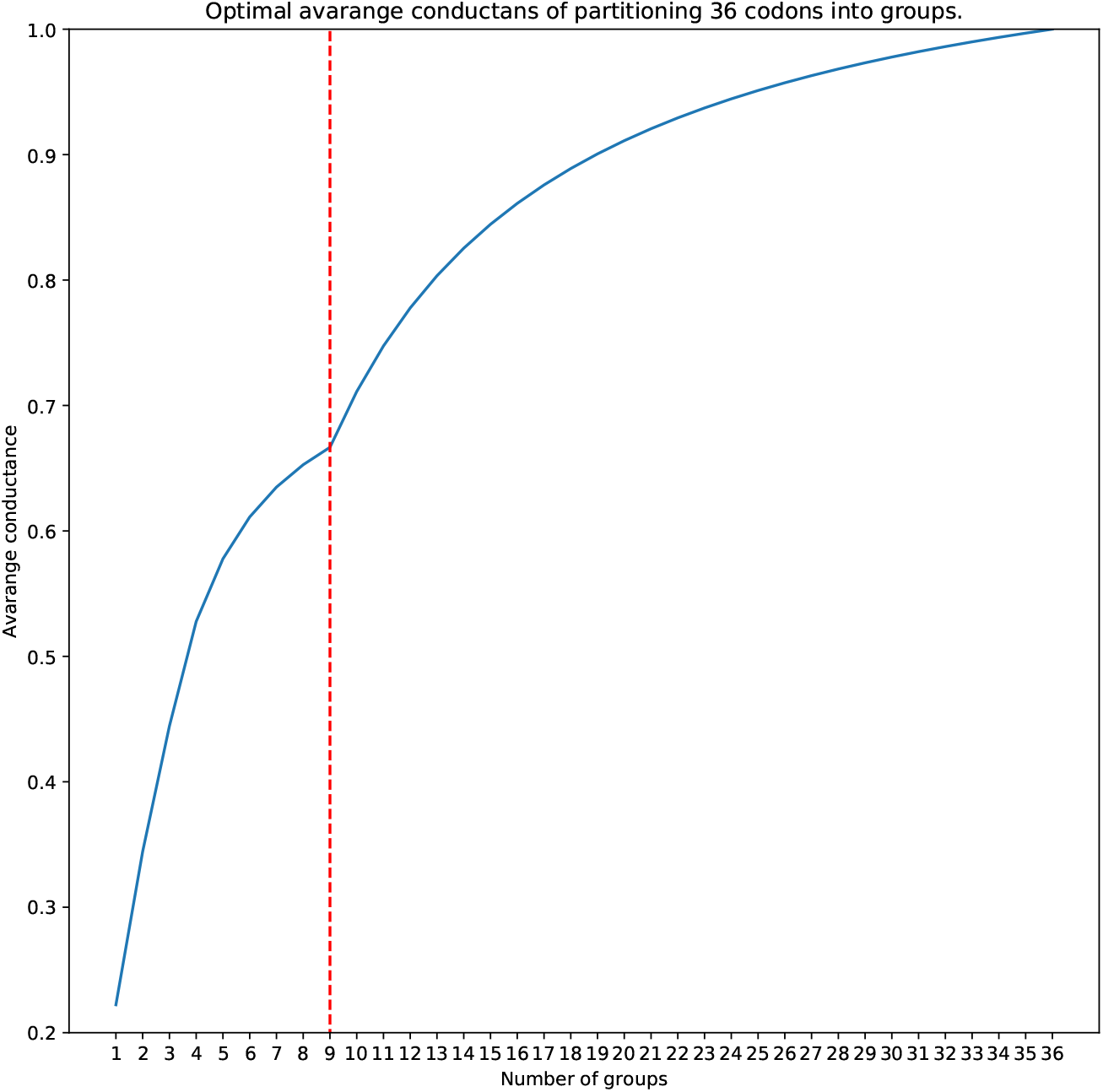
The minimum of the average set conductance 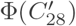 (blue line) in relation to the number *n* of potential codon blocks which would encode new genetic information. The minimum of Φ was found over all possible partitions of the set 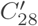 containing 36 vacant codons in exactly *n* disjoint codon blocks. The red dashed line shows the minimum of the average set conductance calculated for *n* = 9. As we can see, *n* = 9 is a deflection point, in which the rate of the curve increase is changing.

### 4.3 The balanced extended genetic code

As mentioned in the previous section, we can describe the optimal genetic code structure, in terms of the minimum of 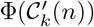, which encode ncAAs for the respective *n* and *k*. However, according to the representation 2, the extended genetic code is in fact composed of 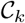, which encodes the canonical information and 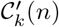, which encodes exactly *n* ncAAs. Following the definition of 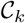, we get that the number of connections between these two parts of the code is as small as possible, which may causes a low probability of potential reversion between the new and old information. It is very useful from experimental point of view, when we want to keep the information about the canonical amino acids and the stop translation, and simultaneously not lose the new information encoded in the vacant codons. However, this property does not include any information about properties of codon groups. On the other hand, the average code conductance 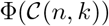 calculated for the whole system give us only a general overview of the quality of codon blocks. In this context, the balance measure *B* appears to be especially useful in studying properties of codon groups belonging to 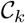 and 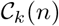. Thanks to that, we can compare the quality of coding system for the new and old information.

In this work, we tested the balance in the case when 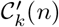 attained lower bound of 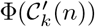. Figure 5 presents the balance values *B* calculated for the respective *k* and *n*. As we can see, the extended genetic code is extremely unbalanced for small *n*, i.e. *B* < 1, which indicates that 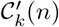 have in general a better codon blocks structure in comparison to respective 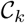 in terms of the average conductance. In all considered cases *B* increases with the number of newly incorporated ncAAs. Moreover, in some cases, it is possible to find a balanced codes for which *B* are around one. What is more, the number of newly included ncAAs, required to obtain the balanced code, is in some cases quite large. For example, in the case of *k* = 28, possible balanced genetic codes are obtained for *n* = 28, 29, 30. This result shows in fact a huge redundancy level of the standard genetic code.

**Figure 5.**
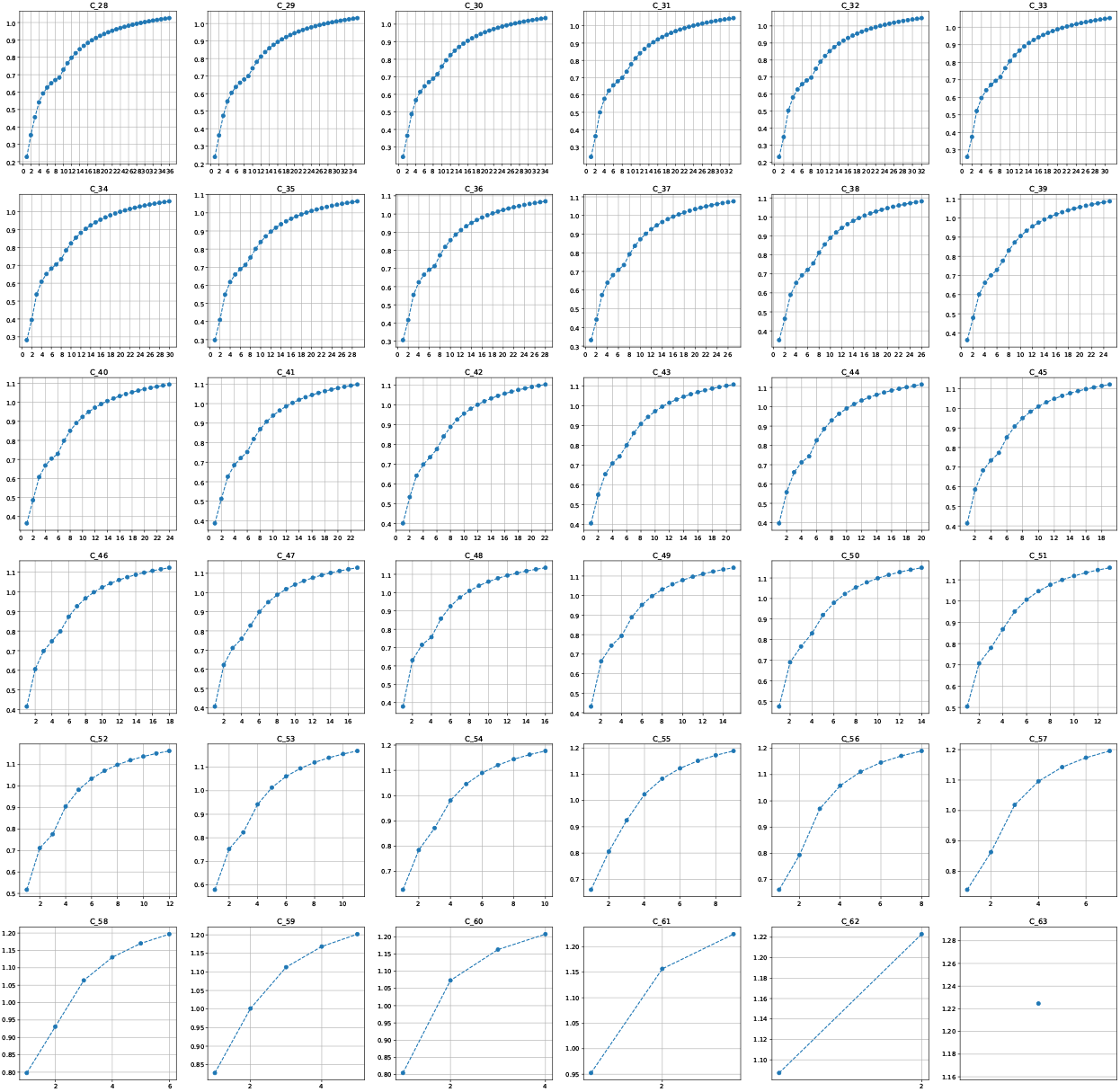
The balance *B* calculated for respective *k* and *n* under the assumption that 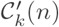 attains the lower bound of 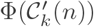. As we can see, the extended genetic code is in general strongly unbalanced for small *n* i.e. the number of newly incorporated ncAAs, in comparison to the code encoding 20 canonical amino acids and one stop translation signal.

## 5 Conclusions

The redundancy of the standard genetic code suggests that this coding system can be extended. In literature, we can find several approaches to this problem. These findings encouraged us to start studying the issue of the optimal extension of the standard genetic code from theoretical perspective. In this paper, we proposed a method of genetic code extension using graph theory approach. Following this methodology, we described the smallest set of codons still encoding 21 canonical items (20 amino acids with one stop translation signal) and characterizing by the minimal set conductance for its size. This property provides the smallest number of connections between codons in the restricted code and the set of vacant codons, which can be assigned to new genetic information. Thanks to that, we minimized the possibility of reversion between these two parts of the code. What is more, we investigated the optimal structure of codon blocks that encode new information and finally we found the lower bounds for the optimal structure of codon blocks assigned to potential ncAAs. In addition, the introduced balance measure allows us for finding extended genetic codes balanced in terms of the average conductance.

It should be noted that the results presented here require some theoretical assumptions. First of all, we proposed a general approach, which does not take into account properties of coded amino acids or other compounds associated with the newly added codons. Secondly, we need exactly 64 types of tRNAs, which can be used to decode unambiguously respective codons. Nevertheless, it seems reasonable to investigate the problem of possible extensions of the standard genetic code starting from the general foundations. Interestingly, using these assumptions, we found several interesting limitations on the number of codons required to encode canonical information and also on the codon blocks which would encode new information.

## References

Anderson et al., 2004. Anderson, J. C., Wu, N., Santoro, S. W., Lakshman, V., King, D. S., and Schultz, P. G. (2004). An expanded genetic code with a functional quadruplet codon. Proc Natl Acad Sci USA, 101(20):7566–7571.

Błażej et al., 2018a. Błażej, P., Kowalski, D., Mackiewicz, D., Wnetrzak, M., Aloqalaa, D., and Mackiewicz, P. (2018a). The structure of the genetic code as an optimal graph clustering problem. https://www.biorxiv.org/content/early/2018/05/28/332478.

Błażej et al., 2019a. Błażej, P., Wnetrzak, M., Mackiewicz, D., Gagat, P., and Mackiewicz, P. (2019a). Many alternative and theoretical genetic codes are more robust to amino acid replacements than the standard genetic code. Journal of Theoretical Biology, 464:21–32.

Błażej et al., 2018b. Błażej, P., Wnetrzak, M., Mackiewicz, D., and Mackiewicz, P. (2018b). Optimization of the standard genetic code according to three codon positions using an evolutionary algorithm. PLoS One, 13(8):e0201715.

Błażej et al., 2019b. Błażej, P., Wnetrzak, M., Mackiewicz, D., and Mackiewicz, P. (2019b). The influence of different types of translational inaccuracies on the genetic code structure. BMC Bioinformatics, 20(1):114.

Błażej et al., 2020. Błażej, P., Wnetrzak, M., Mackiewicz, D., and Mackiewicz, P. (2020). Basic principles of the genetic code extension. Royal Society Open Science, 7(2):191384.

Błażej et al., 2016. Błażej, P., Wnetrzak, M., and Mackiewicz, P. (2016). The role of crossover operator in evolutionary-based approach to the problem of genetic code optimization. BioSystems, 150:61–72.

Błażej et al., 2018c. Błażej, P., Wnetrzak, M., and Mackiewicz, P. (2018c). The importance of changes observed in the alternative genetic codes. Proceedings of the 11th International Joint Conference on Biomedical Engineering Systems and Technologies (BIOSTEC 2018) - Volume 3: BIOINFORMATICS, pages 154–159.

Bezrukov, 1999. Bezrukov, S. L. (1999). Edge isoperimetic problems on graphs., volume 7, pages 157–197. Akademia Kiado, Budapest.

Chin, 2014. Chin, J. W. (2014). Expanding and reprogramming the genetic code of cells and animals. Annu Rev Biochem, 83:379–408.

Chin, 2017. Chin, J. W. (2017). Expanding and reprogramming the genetic code. Nature, 550(7674):53–60.

Dien et al., 2018. Dien, V. T., Morris, S. E., Karadeema, R. J., and Romesberg, F. E. (2018). Expansion of the genetic code via expansion of the genetic alphabet. Curr Opin Chem Biol, 46:196–202.

Epstein, 1966. Epstein, C. J. (1966). Role of the amino-acid “code” and of selection for conformation in the evolution of proteins. Nature, 210(5031):25–8.

Freeland and Hurst, 1998. Freeland, S. J. and Hurst, L. D. (1998). The genetic code is one in a million. Journal of Molecular Evolution, 47(3):238–248.

Freeland et al., 2000. Freeland, S. J., Knight, R. D., and Landweber, L. F. (2000). Measuring adaptation within the genetic code. Trends Biochem Sci, 25(2):44–5.

Gilis et al., 2001. Gilis, D., Massar, S., Cerf, N. J., and Rooman, M. (2001). Optimality of the genetic code with respect to protein stability and amino-acid frequencies. Genome Biol, 2(11):research0049.1-0049.12.

Goldberg and Wittes, 1966. Goldberg, A. L. and Wittes, R. E. (1966). Genetic code: aspects of organization. Science, 153(3734):420–4.

Haig and Hurst, 1991. Haig, D. and Hurst, L. D. (1991). A quantitative measure of error minimization in the genetic code. Journal of Molecular Evolution, 33(5):412–417.

Hamashima et al., 2018. Hamashima, K., Kimoto, M., and Hirao, I. (2018). Creation of unnatural base pairs for genetic alphabet expansion toward synthetic xenobiology. Curr Opin Chem Biol, 46:108–114.

Hohsaka et al., 1996. Hohsaka, T., Ashizuka, Y., Murakami, H., and Sisido, M. (1996). Incorporation of nonnatural amino acids into streptavidin through in vitro frame-shift suppression. J Am Chem Soc, 118(40):9778–9779.

Ishikawa et al., 2000. Ishikawa, M., Hirao, I., and Yokoyama, S. (2000). Synthesis of 3-(2-deoxy-beta-d-ribofuranosyl)pyridin-2-one and 2-amino-6-(n,n-dimethylamino)-9-(2-deoxy-beta-d-ribofuranosyl)purine derivatives for an unnatural base pair. Tetrahedron Letters, 41(20):3931–3934.

Italia et al., 2017. Italia, J. S., Addy, P. S., Wrobel, C. J., Crawford, L. A., Lajoie, M. J., Zheng, Y., and Chatterjee, A. (2017). An orthogonalized platform for genetic code expansion in both bacteria and eukaryotes. Nat Chem Biol, 13(4):446–450.

Iwane et al., 2016. Iwane, Y., Hitomi, A., Murakami, H., Katoh, T., Goto, Y., and Suga, H. (2016). Expanding the amino acid repertoire of ribosomal polypeptide synthesis via the artificial division of codon boxes. Nature Chemistry, 8(4):317–325.

Kimoto et al., 2009. Kimoto, M., Kawai, R., Mitsui, T., Yokoyama, S., and Hirao, I. (2009). An unnatural base pair system for efficient pcr amplification and functionalization of dna molecules. Nucleic Acids Res, 37(2):e14.

Malyshev et al., 2009. Malyshev, D. A., Seo, Y. J., Ordoukhanian, P., and Romesberg, F. E. (2009). Pcr with an expanded genetic alphabet. J Am Chem Soc, 131(41):14620–1.

Massey, 2008. Massey, S. E. (2008). A neutral origin for error minimization in the genetic code. Journal of Molecular Evolution, 67(5):510–516.

Neumann et al., 2010. Neumann, H., Wang, K., Davis, L., Garcia-Alai, M., and Chin, J. W. (2010). Encoding multiple unnatural amino acids via evolution of a quadruplet-decoding ribosome. Nature, 464(7287):441–4.

Noren et al., 1989. Noren, C. J., Anthony-Cahill, S. J., Griffith, M. C., and Schultz, P. G. (1989). A general method for site-specific incorporation of unnatural amino acids into proteins. Science, 244(4901):182–8.

Novozhilov et al., 2007. Novozhilov, A. S., Wolf, Y. I., and Koonin, E. V. (2007). Evolution of the genetic code: partial optimization of a random code for robustness to translation error in a rugged fitness landscape. Biol Direct, 2:24.

Ohtsuki et al., 2001. Ohtsuki, T., Kimoto, M., Ishikawa, M., Mitsui, T., Hirao, I., and Yokoyama, S. (2001). Unnatural base pairs for specific transcription. Proc Natl Acad Sci USA, 98(9):4922–4925.

Ozer et al., 2017. Ozer, E., Chemla, Y., Schlesinger, O., Aviram, H. Y., Riven, I., Haran, G., and Alfonta, L. (2017). In vitro suppression of two different stop codons. Biotechnol Bioeng, 114(5):1065–1073.

Plotkin and Kudla, 2011. Plotkin, J. B. and Kudla, G. (2011). Synonymous but not the same: the causes and consequences of codon bias. Nature Reviews Genetics, 12(1):32–42.

Santos and Monteagudo, 2017. Santos, J. and Monteagudo, A. (2017). Inclusion of the fitness sharing technique in an evolutionary algorithm to analyze the fitness landscape of the genetic code adaptability. BMC Bioinformatics, 18(1):195.

Santos et al., 2011. Santos, M. A. S., Gomes, A. C., Santos, M. C., Carreto, L. C., and Moura, G. R. (2011). The genetic code of the fungal ctg clade. Comptes Rendus Biologies, 334(8-9):607–611.

Sonneborn, 1965. Sonneborn, T. (1965). Degeneracy of the genetic code: extent, nature, and genetic implications., pages 377–397. Academic Press, New York.

Wnetrzak et al., 2018. Wnetrzak, M., Blaizej, P., Mackiewicz, D., and Mackiewicz, P. (2018). The optimality of the standard genetic code assessed by an eight-objective evolutionary algorithm. BMC Evolutionary Biology, 18:192.

Wnetrzak et al., 2019. Wnetrzak, M., Błażej, P., and Mackiewicz, P. (2019). Optimization of the standard genetic code in terms of two mutation types: Point mutations and frameshifts. BioSystems, (181):44–50.

Woese, 1965. Woese, C. R. (1965). On the evolution of the genetic code. Proc Natl Acad Sci USA, 54(6):1546–52.

Yang et al., 2007. Yang, Z., Sismour, A. M., Sheng, P., Puskar, N. L., and Benner, S. A. (2007). Enzymatic incorporation of a third nucleobase pair. Nucleic Acids Res, 35(13):4238–49.

Young and Schultz, 2018. Young, D. D. and Schultz, P. G. (2018). Playing with the molecules of life. ACS Chem Biol, 13(4):854–870.

